# Emo-FilM: A multimodal dataset for affective neuroscience using naturalistic stimuli

**DOI:** 10.1101/2024.02.26.582043

**Authors:** Elenor Morgenroth, Stefano Moia, Laura Vilaclara, Raphael Fournier, Michal Muszynski, Maria Ploumitsakou, Marina Almató-Bellavista, Patrik Vuilleumier, Dimitri Van De Ville

## Abstract

The extensive Emo-FilM dataset stands for **Emo**tion research using **Fil**ms and f**M**RI in healthy participants. This dataset includes detailed emotion annotations by 44 raters for 14 short films with a combined duration of over 2½ hours, as well as recordings of respiration, heart rate, and functional magnetic resonance imaging (fMRI) from a different sample of 30 individuals watching the same films. The detailed annotations of experienced emotion evaluated 50 items including ratings of discrete emotions and emotion components from the domains of appraisal, motivation, motor expression, physiological response, and feeling. Quality assessment for the behavioural data shows a mean inter-rater agreement of 0.38. The parallel fMRI data was acquired at 3 Tesla in four sessions, accompanied with a high-resolution structural (T1) and resting state fMRI scans for each participant. Physiological recordings during fMRI included heart rate, respiration, and electrodermal activity (EDA). Quality assessment indicators confirm acceptable quality of the MRI data. This dataset is designed, but not limited, to studying the dynamic neural processes involved in emotion experience. A particular strength of this data is the high temporal resolution of behavioural annotations, as well as the inclusion of a validation study in the fMRI sample. This high-quality behavioural data in combination with continuous physiological and MRI measurements makes this dataset a treasure trove for researching human emotion in response to naturalistic stimulation in a multimodal framework.

## 1. Background and Summary

Neuroimaging under naturalistic conditions is a growing field within neuroscience, which has been proven useful in a variety of subjects including language [1], [2], social cognition [3] and emotion [4], [5]. While the nature of functional Magnetic Resonance Imaging (fMRI) intrinsically limits the observation of naturalistic conditions, movies and films can be easily implemented as naturalistic paradigms in the scanner. In particular, film fMRI opens a range of new pathways for understanding the brain, as reflected by an increasing push towards naturalistic and other non-traditional paradigms in the field [6].

Film fMRI is especially promising for emotion research, as films evoke a higher intensity of emotion compared to other methods [7], [8]. The ecological validity of the participants’ emotion experience is also superior when elicited by films, because events dynamically unfold over time and allow a natural evolution of emotions across successive moments. There is an increasing amount of publicly available fMRI datasets that include film watching (e.g. [3], [9],[10], [11]), yet without annotations regarding emotion elicitation these have limited value for affective neuroscience. Acquiring such annotations is very resourceful as to date it is not possible to reliably extract rich information on experienced emotion in an automated fashion from audiovisual contents themselves, e.g., by relying upon artificial intelligence. Therefore, a large community effort is needed to produce and share film fMRI datasets that include detailed emotion annotations. Furthermore, the inclusion of detailed physiological measurements is needed to better understand the sources of the neural signal and the effects of emotional stimuli on emotion [12].

While neuroscience research on emotion has long been dominated by bidimensional (valence and arousal) or core affect models [13], appraisal theories are currently receiving increasing attention and already made important contributions in psychology [14], [15]. These theories comprise a group of models predicting that emotions are determined by an individual’s appraisal of a current event or stimuli in relation to their goals and needs [15]. Although appraisals constitute a well-established mechanism of emotion elicitation, this framework was only rarely investigated in neuroimaging studies [16], [17]. There is a need to better characterise the neural processes of cognitive appraisal and subsequent emotional responses, especially given the potential to inform our understanding of perturbed emotion processing in psychopathology by directly building on these insights.

Here we focus on a specific appraisal theory, the Component Process Model (CPM), initially proposed by Scherer [18]. The CPM describes how emotion is composed of a set of five distinct components: appraisal, motivation, expression, physiology, and subjective feeling. Compared to other frameworks, the CPM comes with a larger library of resources for empirical research. Most notably, the Geneva Emotion Recognition tool (GRID) instrument provides a collection of emotion words and features in accordance with the CPM and other theoretical approaches to emotion, such as the dimensional and the basic emotion approach [19]. A small number of fMRI studies have based their investigations on the CPM so far [16], [20], [21], however rating data (available to the wider community) did not include rich moment-by-moment annotations of film content but were based on preselected snapshots or experimentally induced [16], [20], [21],

We present the EmoFilm dataset, that was obtained by combining an annotation with a neuroimaging study. In the annotation study part, we use a selection of 16 short films for which valence and arousal ratings are already available elsewhere as part of the LIRIS-ACCEDE database [22] and for aesthetic highlight [23]. We added to these existing data new continuous annotations for another 55 emotion-relevant items, 13 from the domain of discrete emotions and 42 from the categories of appraisal, motivation, expression, physiology, and subjective feeling based on the GRID instrument [19]. Based on ratings from 44 annotators, we calculated a consensus annotation to describe the general pattern of behavioural responses to the films’ content. The data for this part of our study has been organised in BIDS format and can be found on OpenNeuro [24] (doi:10.18112/openneuro.ds004872.v1.0.0). In the neuroimaging study part, we collected fMRI and physiological data from an independent sample of 30 participants watching 14 of these short films where reliable consensus ratings could be calculated in the annotation study. We also included a behavioural task after fMRI scanning during which participants rated short clips taken from the same films in order to validate the continuous ratings obtained in the annotation study. The data for this part of our study has been organised in BIDS format and can be found on OpenNeuro [25] (doi:10.18112/openneuro.ds004892.v1.0.0).

## 2. Annotation Study

### 2.1. Methods

#### 2.1.1. Participants

Forty-four participants (23 female) were recruited over the course of the study to perform movie annotations remotely using their own computers. The mean age was 25.31 with a range from 20 - 39. Inclusion criteria were high oral comprehension level for English, no history of psychiatric or neurological diseases, no recreational drug use as well as no current neuropharmacological medication. Despite the online nature of data collection, we deliberately recruited participants locally, from Geneva university and the surrounding population, expecting a higher data quality with stronger motivation and better match with subsequent fMRI sample. Recruitment was performed via a questionnaire that was circulated online in relevant groups and forums within the university community and the wider population in the Geneva area. As four participants eventually failed to complete the whole experiment, we recruited four additional participants to compensate for missing data. Participants were reimbursed with 20 CHF/hour upon completion of the experiment. In total, forty-four participants completed the experiment between November 2020 and February 2021. One participant completed the experiment in January 2022 and another in October 2022, after they were recruited to replace corrupt or missing data. Ethical approval was given by the Geneva Cantonal Commission for Ethics in Research (protocol No 2018-02006). The study complied with the Code of Human Research Ethics (2014). All participants gave written informed consent prior to taking part in the study and were transparently informed of research goals.

#### 2.1.2. Materials

##### 2.1.2.1. Films

Emotion annotations were acquired for 16 short films taken from the films included in the LIRIS database [22] and previously used for affective research. All selected films are free to share under Creative Commons licences. They were chosen based on their potential to evoke a broad range of emotions, but also based on logistical considerations, including film duration, diversity of content or format. For the purpose of this research, the beginning and end credits were cut. Our dataset includes the resulting clips ranged in duration from 6 minutes 42 seconds to 17 minutes and 8 seconds (average 11 minutes 47 seconds). Table 1 details the duration of each film with information on their genre and content.

**Table 1.** List of films used in the fMRI study, with film duration, scan duration, content description, film genre, and average ratings of absorption, enjoyment, and interest given by participants after scanning. * indicates films that were not included in the fMRI acquisition.

| Film | Film duration (s) | Scan duration (TRs) includes washout | Description | Genre | Absorption | Enjoyment | Interest |
| --- | --- | --- | --- | --- | --- | --- | --- |
|  |  |  |  |  | Mean of ratings on a scale 0-100 |  |  |
| After The Rain | 8:16 | 534 | A man contemplates the meaning of life and human interaction on a rainy day in the city. | Drama | 49 | 47 | 49 |
| Between Viewings | 13:28 | 776 | A disillusioned estate agent is forced to re-examine his life when asked to sell his childhood home | Comedy/ Drama | 64 | 69 | 68 |
| Big Buck Bunny | 8:10 | 528 | An enormous, fluffy, and adorable rabbit is harassed by a bullying gang of squirrels | Animation / Comedy | 74 | 78 | 68 |
| Chatter | 6:45 | 464 | A girl witnesses a horrible sight online, then the electricity is cut off inside her apartment. When the light returns, she feels that she is not alone. | Thriller | 69 | 50 | 60 |
| Damaged Kung Fu* | 15:22 | n/a | Two friends are shooting a Kung Fu film together, when one quits thinking his friend is having an affair with his girl friend. The crew try to replace him in the shoot with little success. | Action/ Comedy | n/a | n/a | n/a |
| First Bite | 9:59 | 613 | A teenage girl is discovering the power of seduction. | Romance | 55 | 51 | 52 |
| Lesson Learned | 11:07 | 665 | A young man turns his life around after being involved in gang violence. | Drama | 60 | 53 | 61 |
| Payload | 16:48 | 928 | A man must sacrifice everything to save his family. | Drama/ Sci-Fi | 55 | 53 | 60 |
| Riding the Rails* | 13:14 | n/a | A boy's grandfather tries to cheer him up on his birthday with a new train for his train set. | Drama | n/a | n/a | n/a |
| Sintel | 12:02 | 710 | A young woman embarks on a dangerous quest to find her lost friend, a dragon. | Animation / Fantasy | 85 | 80 | 83 |
| Spaceman | 13:25 | 744 | A young man sets out on a very curious and unique path to realise his dream of being an astronaut. | Romance/ Drama | 60 | 65 | 68 |
| Superhero | 17:08 | 945 | A single mother cares for her son with terminal cancer. | Drama | 65 | 56 | 60 |
| Tears Of Steel | 9:48 | 607 | A group of soldiers and scientists try to stop an army of robots that | Action/ | 68 | 69 | 71 |
|  |  |  | threatens the planet by correcting a past mistake. | Sci-Fi |  |  |  |
| The Secret Number | 13:04 | 757 | A psychiatrist is compelled by his patient, an obsessive mathematician, to consider the existence of a secret integer between three and four. | Drama | 77 | 80 | 81 |
| To Claire From Sonny | 6:42 | 460 | A young man writes a letter to his first true love. | Drama | 71 | 73 | 73 |
| You Again | 13:18 | 768 | A chance encounter between two former high school sweethearts forces them to face the ways they have - and have not - changed. | Romance/<br>Drama | 45 | 49 | 46 |

##### 2.1.2.2. Annotation Items

In our study, 55 items were annotated comprising 42 items from the categories of Appraisal, Expression, Physiology, Motivation and Feeling that were adapted from the CoreGRID instrument [19], plus a further 13 terms for discrete emotions (see Supplementary table 1 for a list and description of all items).

##### 2.1.2.3. Questionnaires

A number of questionnaires were given upon completion of the emotion annotation task. We used total scores computed from these questionnaires. The Depression Anxiety Stress Scales (DASS; [26]) was used to assess affective state over the previous seven days. We also employed an in-house scale to gauge how people were affected by the Covid-19 pandemic and its consequences. The scale includes items rating the pandemics’ effect on social support, mental health, concerns about getting infected or infecting others, worries about the future and impact on cognitive function (internal consistency; alpha = 0.80). The BIS/BAS Scale was used to measure the behavioural approach system on the subscales drive, fun seeking, and reward responsiveness, and the behavioural inhibition system [27]. We also administered the Emotion Regulation Questionnaire (ERQ), which probes two facets of emotion regulation: Cognitive Reappraisal and Expressive Suppression [28], as well as the Big Five Inventory (BFI, [29]) which was used to create scores of Extraversion, Agreeableness, Conscientiousness, Openness, and Neuroticism. Description of the sample based on these questionnaires can be found in Supplementary table 2.

#### 2.1.3. Procedure

Upon recruitment, participants were contacted with detailed information about the study and invited for a video call with a researcher. During this meeting, participants were instructed how to download and use the annotation software, how to access their annotation tasks, and how to upload completed annotations onto a secure online platform. Participants were further supplied with detailed instructions pertaining to the interpretation and directionality of the items they were assigned to ensure uniform interpretations across our sample. In addition, they were given brief descriptions of the items they rated in written form, such that they could consult them when needed. All materials were accessed by participants via this dedicated online platform.

Participants were instructed to complete a list of annotation tasks in a given order at their own pace. This list was prepared for each participant and consisted of all films with six rating items each, resulting in 96 film by item combinations, sorted in blocks by item. This means that participants would annotate one item for all 16 films, before moving on to the next item. The films in each block were given in random order. We do not expect adverse effects due to participants viewing each film multiple times based on the relatively long delay between repetitions and findings that repeating a specific emotional stimulus has only a negligible effect on self-reported emotional feelings [30].

Participants were encouraged to complete all annotations within six weeks. Upon completion of a session, they were instructed to upload their response files onto a secure system. This ensured that the quality of their annotations could be monitored continuously.

To obtain online ratings, we used an adapted version of the software CARMA [31], specifically developed for film annotations. The main customizations were related to the annotation scale adapting to the current item, which was specified in the file name of each rated film. The sampling rate within CARMA software was fixed to 1Hz. To complete their annotation tasks, participants would load the prepared video files according to their order and then move a mouse-controlled cursor along a bar on the computer screen to continuously annotate an item. Continuous quality control was performed using visual inspection of time-courses and analysis of agreement between raters when applicable. Participants received feedback if their time courses appeared too “synthetic” (e.g., box-shaped or constant) or if there was an unexpectedly high discrepancy between their annotations and the rest of the cohort. No participants were excluded based on annotation quality.

Upon completion of the emotion annotation task, participants completed a number of psychometric questionnaires on an online platform.

#### 2.1.4. Calculation of Consensus Annotation

All annotation time series were z-scored across films within each rater before further processing. Individual missing values in time series were replaced with the mean of the two neighbouring values. Constant time series were discarded and not included in the calculation of the consensus annotation, as were time series with outliers beyond a z value of 15 or -15. Finally, the quality of annotations was assessed using Pearson’s correlation coefficient (r). Specifically, for each item and each film, r was calculated between each pair of annotations across participants (resulting in 6 r values), and then averaged across pairs to result in one value of agreement per movie and item. If the inclusion of a time series reduced mean r between all raters by more than .20, the time series was discarded for calculation of a consensus annotation. Finally, in a few exceptional cases where five annotations were available (due to additional recruitment, see above), we removed the annotation with the lowest average correlation with the other annotations. The remaining complete time series were averaged per item and per film to form the consensus annotation. Each consensus annotation time series was based on the average of at least three raters.

### 2.2. Annotation Quality

Annotation quality was also assessed for each item and each film pairing using Pearson correlation. Agreement was strongly dependent on the item that was annotated with the highest agreement being r = 0.58 for *PleasantOther* and the lowest being r = -0.01 for *Jaw*. Mean agreement also differed between films (ranging from 0.21-0.50). While it is natural for agreement to vary between items and films, we recommend that items with mean agreement of smaller r = 0.20 cannot be considered reliable. Consequently, the following items were removed from all further analysis as their agreement across all movies was smaller than r = 0.10: *Breathing* (r = 0.14), *Consequences* (r = 0.12), *Movement* (r = 0.07), *EyesOpen* (r = 0.06), *Jaw* (-0.01). We also removed two films that did not achieve a mean agreement of at least 0.25 across all items: *Riding the Rails* (r = 0.21) and *Damaged Kung Fu* (r = 0.245). From the remaining 14 films and 50 items, 2853 individual annotation time series were available. Of these, 27 annotations were removed because of constant segments, and 23 because of outliers beyond a z-value of 15 or -15. In addition, 125 time series were removed as being deviant (their exclusion improved mean inter-rater agreement by more than r = 0.2) and 21 annotations were removed as they were the worst of five in terms of inter-rater agreement.

Consequently, Figure 1 shows a detailed summary of agreement overall (A), within films (B), and across rating items (C) and CPM categories (D), after removal of the two films and five items with poor reliability. The final dataset therefore includes annotations from 50 items for 14 films, with an average agreement for films across items between r = 0.28 - 0.54, and agreement for items across films between r = 0.21 - 0.61. Of the 700 film and item combinations in these data, the final consensus annotation was calculated based on three annotations for 143 cases, and based on four annotations for all remaining ones. The mean agreement across all items and films was r = 0.38 (see Supplementary Table 3 for detailed Inter-rater agreement across all films and items).

**Figure 1.**
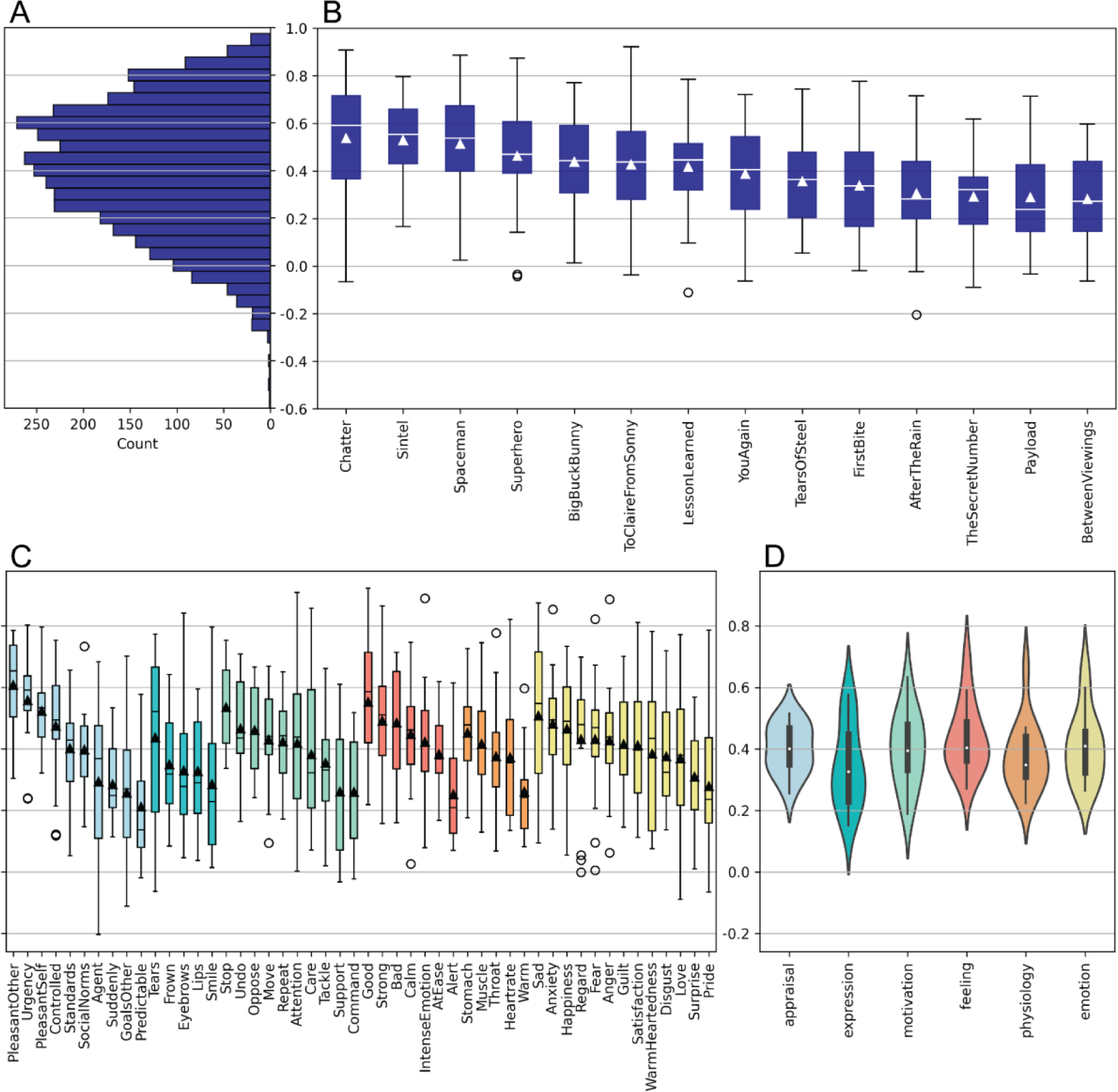
Inter-rater agreement in annotation study. (A) Histogram of inter-rater agreement between all valid pairs of ratings. (B) Distribution of inter-rater agreement by film based on all valid pairs of ratings. (C) Distribution of inter-rater agreement by item based on all valid pairs of ratings, including GRID items and discrete emotion terms. Bars are coloured by components according to the CPM framework, as in (D). (D) Distribution of inter-rater agreement by components and discrete emotions.

We furthermore report the average value from the consensus annotation for each item in each film in Figure 2. This illustrates the expected variety of relative emotion intensity between films, but also shows that various emotion dimensions were generally elicited within a given film.

**Figure 2.**
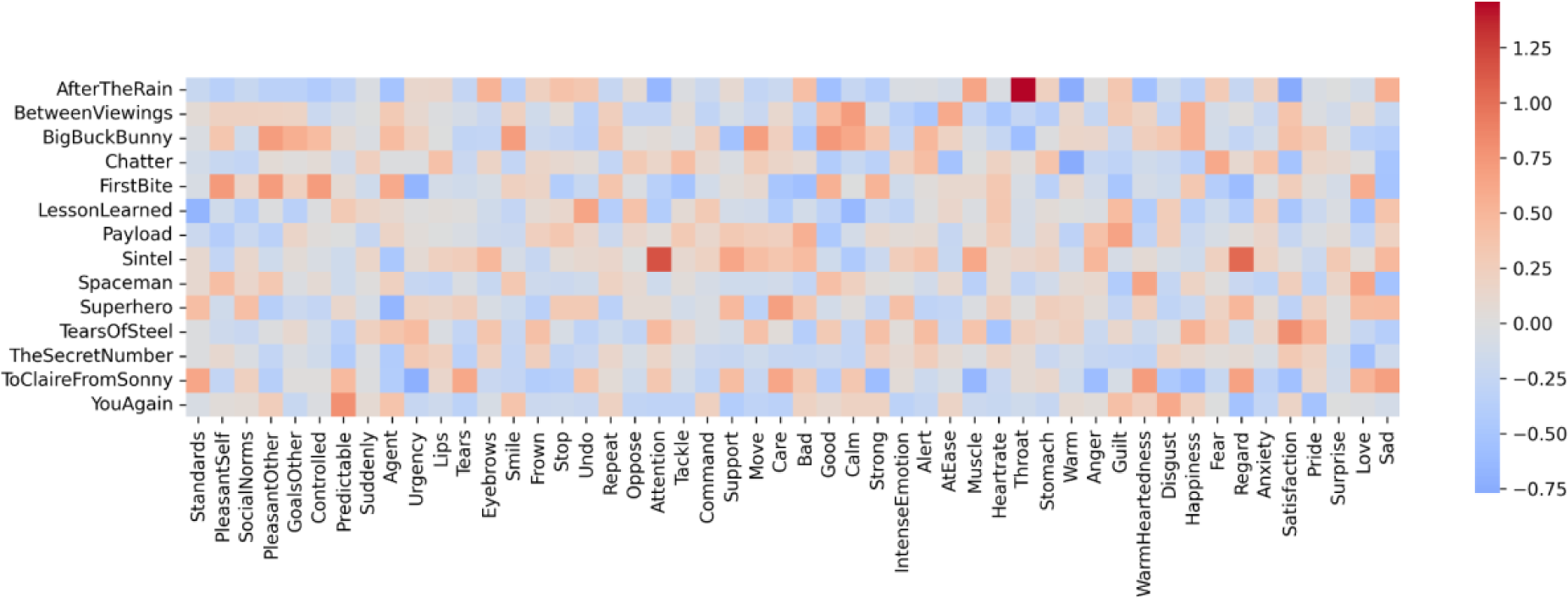
Average value of consensus annotation for each item and film. High values (red) indicate films that were rated consistently higher on this item relative to other films. Conversely, low values (blue) indicate films that were rated consistently lower on this item relative to other films.

## 3. fMRI study

### 3.1. Methods

#### 3.1.1. Participants

Thirty-two healthy volunteers were recruited for the fMRI experiment. Two had to be excluded during the first session, because one could not tolerate lying in the MRI scanner and one had strong artefacts due to dental braces. Consequently, 30 subjects (18 female) completed the fMRI experiment, none of which partook in the annotation study. All subjects were healthy adults between 18 and 35 years old (average 25.83, std = 3.60) and right-handed as confirmed with the Edinburgh Inventory [32]. All had normal or corrected-to-normal vision including full colour vision, high level of English language comprehension, no history of any neurological or psychiatric condition, and none reported using neuropharmacological or recreational drugs. Ethical approval was given by the Geneva Cantonal Commission for Ethics in Research (protocol No 2018-02006). The study complied with the Code of Human Research Ethics (2014). All participants gave written informed consent prior to taking part in the study and were transparently informed of the research goals at all times.

#### 3.1.2. Materials

##### 3.1.2.1. Films

We used 14 short films selected from the previous annotation study (see Table 1). Two films were not included because of unreliable consensus annotations (*Damaged Kung Fu* and *Riding the Rails*). We used the same clips as before, without beginning and end credits. The average duration of these films was 11 minutes 26 seconds.

##### 3.1.2.2. Annotation Items

After scanning, we included an offline behavioural rating phase to validate the annotations obtained from other participants in the previous study. We used a subset of 48 items comprising 34 items from the categories of Appraisal, Expression, Physiology, Motivation and Feeling taken from the CoreGRID [19] and 13 discrete emotion terms. We did not include items for which we found no reliable consensus annotation in the previous study (see Supplementary Table 1 for list and description of all items).

##### 3.1.2.3. Questionnaires

We used the same battery of questionnaires as in the annotation study. A description of the sample based on these questionnaires is provided in Supplementary Table 2.

#### 3.1.3. Procedure

##### 3.1.3.1. Imaging Experiment

The experiment spanned over four fMRI sessions each lasting approximately two hours. During these sessions, subjects watched between two and five short films in the MRI scanner and subsequently rated their emotion experience during watching in the offline behavioural test. Additionally, subjects underwent a 10-minute resting-state scan in the first session during which they were asked to keep their eyes open and fixate a crosshair on the screen. Each subject watched the films in pseudo-random order, distributed over the four sessions. Stimulus presentation was programmed in MATLAB 2012, using the Psychophysics Toolbox extensions [33], [34], [35]. Each film run started and ended with a 90 second washout period during which a crosshair was presented centrally on the screen without auditory stimulation. Between the two washouts, the film was displayed on the screen with the corresponding audio track heard through in-ear plugs. The subjects were instructed to watch the films as they would watch films in their everyday life. At the end of each film, participants responded to three successive questions, displayed in white on a black background on the screen, to indicate their level of absorption (‘I felt absorbed by this movie’), enjoyment (‘I enjoyed this movie’), and interest (‘I thought this movie was interesting’) during film watching. They used a button box to move a slider on the screen up or down a continuous scale to mark their agreement with the respective statement.

##### 3.1.3.2. Validation of Film Annotations

Once outside the MRI scanner, participants completed an offline behavioural task where they rated their emotion experience during the films they had just seen during the fMRI session. This task was programmed in Matlab 2012 using the Psychophysics Toolbox extensions [33], [34], [35]. Participants were given instructions pertaining to the meaning and directionality of the rating items to ensure adequate understanding and uniform interpretations across our sample. During this task, they re-watched selected clips from each film and rated them by moving a slider up and down along a continuous scale (without units or markers) whose extremities indicated high and low experience. Participants rated five different items sequentially after seeing a short clip. In total, each participant watched and rated 292 clips taken from the movies, with an average duration of 7s, equating to 20.52% of the total duration of all films (range = 14.79% - 26.37% of each film). Each item was rated by three to four subjects, such that an average could be computed and compared with annotations observed in the previous study with other participants. To compare the clip ratings with the continuous annotations from the previous study, we applied a linear interpolation over ratings from the clips, then z-scored each time series, and computed an average over all subjects who rated the respective item. This average time series was then compared to the consensus annotation from the behavioural study using Pearson correlation. We also compared the mean inter-rater agreement for each item to the mean correlation between the time courses from this experiment and the consensus annotation for each item.

#### 3.1.4. Physio Acquisition

Participants’ physiological activity was recorded for the whole duration of the fMRI scan with a BIOPAC MP150 monitoring system and recorded with the AcqKnowledge software (version 4.4). Specifically cardiac pulse was collected via photoplethysmogram (BIOPAC TSD200_MRI transducer and PPG100C amplifier), respiratory effort was measured via chest expansion (BIOPAC TSD221-MRI fully pneumatic respiration transducer and RSP100C amplifier), and skin conductance was collected via Electrodermal Activity (EDA) (Cleartrace electrodes 2 RTL and EDA100C amplifier). All signals were sampled at a rate of 1000 Hz. Physiological recordings encompassed the whole acquisition.

#### 3.1.5. Physio Preprocessing

AcqKnowledge proprietary files containing physiological data were organised into the Brain Imaging Data Structure [36] schema with phys2bids [37]. The conversion process simultaneously splits files into runs, keeping extra recording material before and after the run itself (9s on both sides), and converting them into tabular (tsv) format.

After downsampling both cardiac pulse and ventilation data to 40 Hz and applying a low-pass filter (8Hz for cardiac data and 2Hz for ventilation), peaks were detected automatically, with manual supervision, using peakdet [38]. The denoised physiological data was used to model physiological noise with phys2denoise [39], in the form of Heartbeat Interval (HBI) and Respiratory Variance (RV). Briefly, HBI was computed as the median of peak-to-peak distance within a sliding window of 6s, convolved with the opposite of the cardiac response function. RV was computed as the variance of the signal within a sliding window of 8s, convolved with the respiratory response function.

#### 3.1.6. MRI Data Acquisition

MRI scans were acquired on a 3T Siemens Magnetom TIM Trio scanner (Siemens, Erlangen, Germany) using a 32-channel head coil at the Brain and Behaviour Laboratory at the University of Geneva (BBL). Structural T1 weighted images, used for co-registration, were acquired with a GRAPPA sequence TR/TE/flip angle = 1.9 s/2.27 ms/9°, spatial voxel resolution of 1 mm × 1 mm × 1 mm, in plane resolution of 256 × 256 × 192 sagittal slices, field of view 256 mm, and scanning time of 5:04 min.

All functional images were acquired with the same multi-band frequency protocol with a TR/TE/flip angle = 1.3 s/30 ms/64°, field of view 210 mm × 210 mm, slice thickness of 2.5 mm giving a voxel size of 2.5 mm × 2.5 mm × 2.5 mm and whole brain coverage of 54 interleaved slices. Resting-state runs lasted 10 minutes, totalling 460 volumes. The number of volumes acquired for each film and the duration of each film are detailed in Table 1.

#### 3.1.7. fMRI Preprocessing

MRI DICOM files were organised following the BIDS schema with BIDScoin [40], and simultaneously converted to nifti with dcm2niix [41]. fMRI data processing was conducted using FEAT (FMRI Expert Analysis Tool) Version 6.00, part of FSL (FMRIB’s Software Library, www.fmrib.ox.ac.uk/fsl). Images were coregistered to a high-resolution structural, standard space and to the first functional volume of each subject using FLIRT [42], [43]. The following preprocessing pipeline was applied; motion correction using MCFLIRT [43], non-brain removal using BET [44], spatial smoothing using a Gaussian kernel of FWHM 6.0 mm; grand-mean intensity normalisation of the entire 4D dataset by a single multiplicative factor; high pass temporal filtering (Gaussian-weighted least-squares straight line fitting, with sigma = 50 s). We further used FAST segmentation [45] to identify tissue classes at subject level and regress white matter (WM) and cerebrospinal fluid (CSF) from the data together with the six motion regressors derived from image realignment. Finally, we applied defacing to the structural images using pydeface (v. 2.0.0) [42].

### 3.2. Data Quality & Validation

#### 3.2.1. Validation of Annotations

The validity of ratings acquired in the fMRI study was verified by comparing them to the consensus annotation obtained in the preceding annotation study part by computing Pearson correlations between the average ratings from the former and the consensus annotation from the latter. The mean correlation across all films and items was 0.41. This is comparable with the mean inter-rater agreement reported previously. Figure 3 shows histograms of the correlation values between the validation time courses and the consensus annotation, for all combinations between films and items (A) and for the average value within each item (B). The mean agreement with the consensus annotation ranged from 0.08 for *Regard* to 0.70 for *Stop*., but with clear peaks between 0.4 and 0.6.

**Figure 3.**
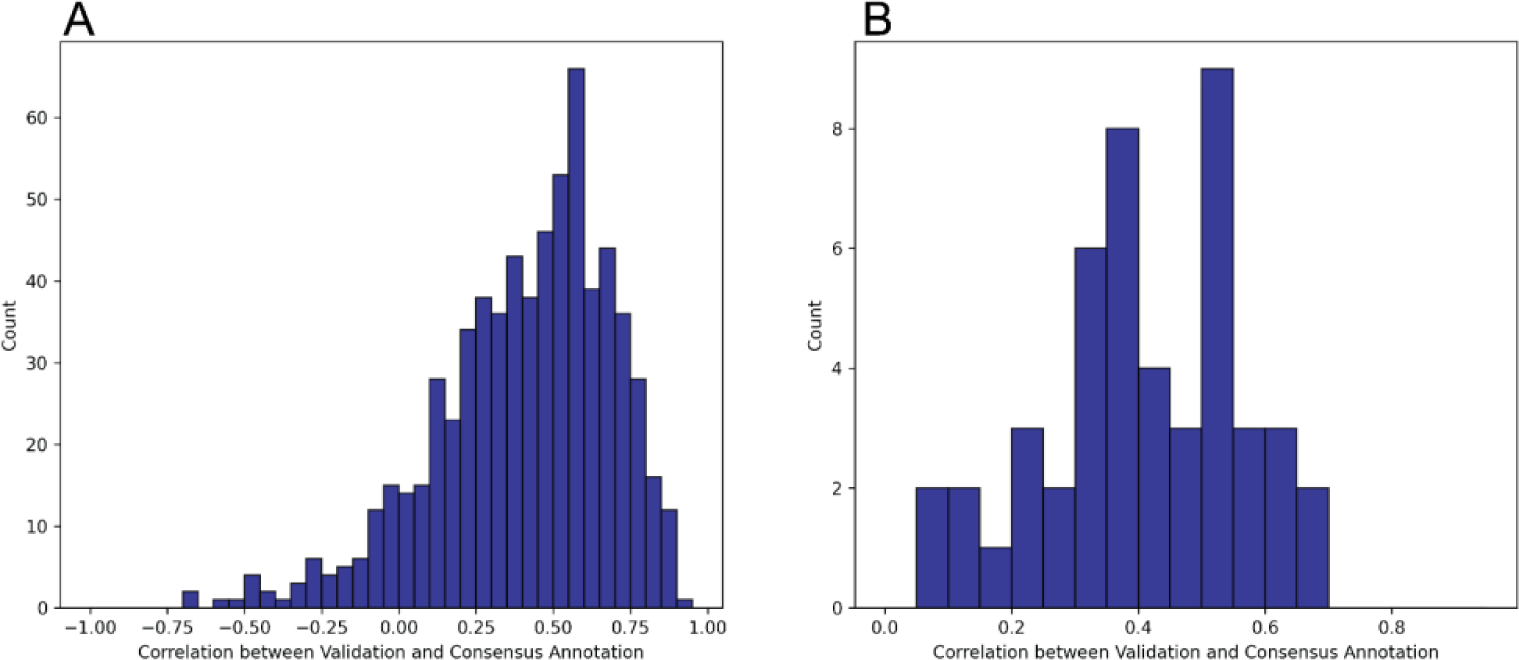
Histograms of the correlation values between validation time courses and the consensus annotation. (A) Agreement for all combinations between films and items and (B) for the average value within each item across films.

For most items, the agreement with the consensus annotation was higher than the inter-rater agreement within individual ratings in the previous study. The correlation between the mean inter-rater agreement for each item in the annotation study and the mean correlation of the validation time course with the consensus annotation was .62. This means that items that reached lower agreement in the annotation study also showed lower agreement between the consensus annotation and the validation time series. This may be a feature of these items, i.e., they may not be experienced as universally as others and potentially depend more on individual differences; or a specificity of the current film material, i.e., some items were not appropriately evoked by the content of selected movies.

#### 3.2.2. fMRI Quality Control

MRIQC (v. 0.16.1) [46] was used to assess quality control of both structural and functional MRI data. Figure 4 and 5 report a subset of the quality metrics computed by MRIQC, for structural and functional volumes respectively. The integral reports can be found in the derivatives of the fMRI dataset on OpenNeuro [25] (doi:10.18112/openneuro.ds004892.v1.0.0)

**Figure 4:**
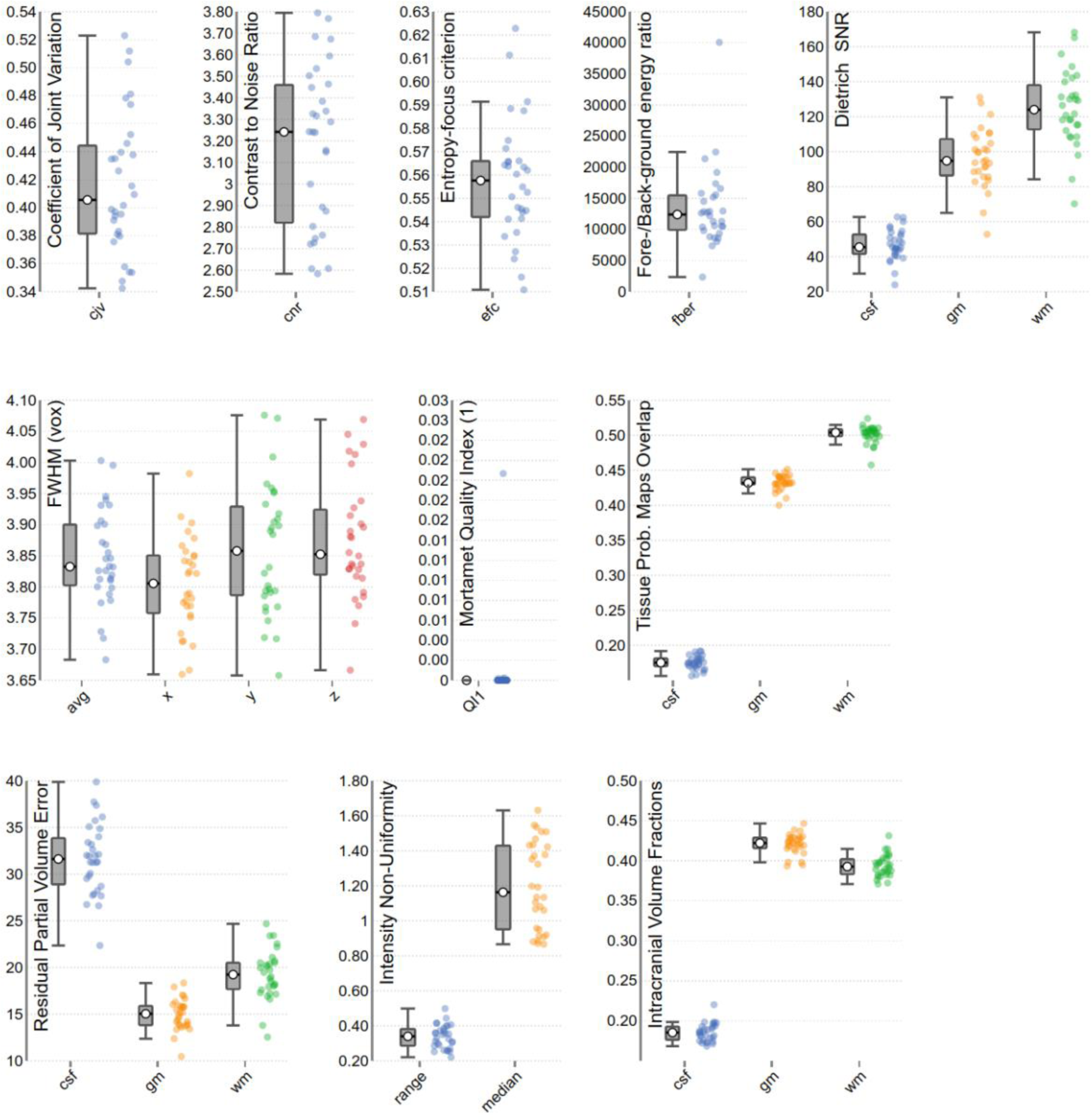
Subgroup of quality metrics of T1w volumes, computed by MRIQC. Each dot represents a volume. For an in-depth explanation of each metric, see [54].

**Figure 5:**
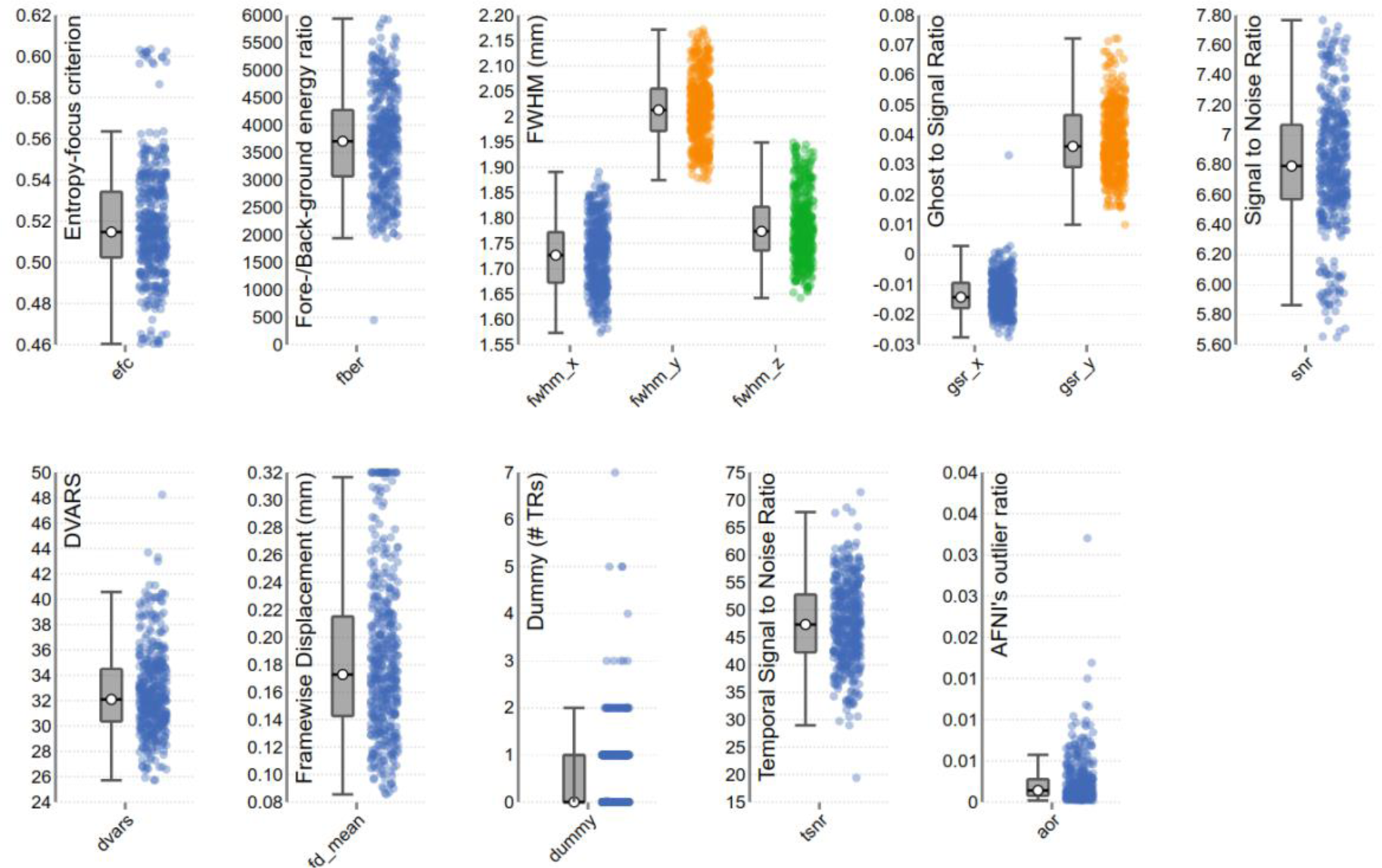
Subgroup of quality metrics of BOLD runs computed by MRIQC. Each dot represents a run. For an in-depth explanation of each metric, see [54].

For the structural images, the coefficient of joint variation [47] indicates absence of heavy head motion, as does framewise displacement (FD) [48] although the intensity non-uniformity index [44] indicates sub-optimal field bias. There seems to be low ghosting and blurring induced by head motion, with few volumes showing an entropy-focus criterion (EFC) higher than 0.58 [49]. The contrast to noise ratio and Dietrich Signal to Noise Ratio (SNR) (a comparison between tissues and background, see [50] are high, especially for Grey Matter (GM) and White Matter (WM), although the average image smoothing median is 3.83 voxels. Mortamet’s Quality Index [51] indicates no voxels with intensity corrupted by artefacts, with the sole exception of a few voxels in the anatomical volume of subject 16.

Most functional volumes show an EFC below 0.57, with the exception of the runs of subject 29, and a film run from subject 20 (The Secret Number). Data smoothness is within the voxel size, the Ghost to Signal Ratio close to 0, although higher in the phase-encoding axis y, and SNR and temporal SNR (measure of MRI signal strength) are high, with very few outliers found by AFNI’s 3dToutcount [52], beside subject 21 rest run, overall indicating acceptable data quality. The number of initial volumes labelled as “dummy”, due to non-steady magnetisation state is within 2 volumes for most runs, with a few exceptions.

Regarding functional runs FD, across all functional runs mean FD was 0.16 mm (SD = 0.10) ranging from 0.09 - 0.55 mm. While FD was generally low, we found a significant difference of FD between film (M = 0.17, SD = 0.10) and rest (M = 0.12, SD = 0.04), with rest having significantly smaller FD (t(448) = 2.72, p < .01). This is contrary to previous findings of reduced motion during film fMRI compared to rest [53]. No subjects were excluded based on MRI image quality.

## 4. Conclusion and Outlook

We present the Emo-FilM dataset, an extensive multimodal corpus for the study of emotion during film watching. This comprises continuous emotion annotations, physiological recordings, and fMRI data for 14 short films that are freely available. This dataset allows tracking emotion differentiation and dynamic unfolding with a high precision and along multiple dimensions. We applied stringent quality control of these data to ensure their suitability to a variety of uses. Regarding the emotion annotations, we demonstrated high levels of inter-rater agreement in the initial annotation study, which was further validated by a behavioural task in an independent sample of participants who performed the subsequent fMRI study. We provide a consensus annotation for 50 different, reliable rating items, based on the average evaluation from at least three annotators for each item, and cover a large dimension space encompassing the five main emotion components (appraisal, motivation, expression, physiology, and feelings) postulated by the CPM framework [18], in addition to 13 traditional discrete emotion terms. Furthermore, we collected fMRI and physiological data in an independent sample of volunteers watching the same short films. This thus extends the subjective annotation measures with objective physiology measures from the peripheral body, including heart rate, electrodermal activity, and respiration recordings, in addition to central neural data acquired with fMRI.

The resulting data from both study parts, i.e., annotations of emotion experience and the corresponding physiological and fMRI recordings, can be put in relation to one another to investigate the effects of emotion experience during film watching in terms of various emotion descriptors. Thus, the Emo FilM dataset constitutes a rich, multimodal tool to investigate cognitive and cerebral processes underlying emotion experience. Although our annotation measures were especially tuned (though not limited) to variables delineated under the theoretical framework of the Component Process Model [18], these may generalise easily to other appraisal models and more generally be integrated with other common emotion theories. The Emo-FilM dataset therefore complements existing annotations of these short films in terms of valence and arousal [22] or aesthetic highlights [23]. While the primary purpose of our new dataset is to reduce the gap between theory in psychology and empirical neuroscience research on emotion, through a refined characterization of brain activity patterns and dynamics in relation to a broad range of emotion experiences, we see many other opportunities offered by these data for a wide variety of research applications.

In addition, given the films used are freely available for research, one can easily extract other relevant features of the same stimuli to further expand the use of this dataset and confront it to multiple theoretical models with various analytical tools. Especially in the context of recent advances in artificial intelligence, automatic extraction of low-level auditory and visual features, language, and semantic information from videos is becoming more accessible and reliable. This information would allow a deeper analysis of the relationship of these features with emotion experience and with changes in physiology and brain activation. An interesting application here may be to better understand how filmmakers use different sensory channels to trigger effective changes in emotion experience. This dataset may also be of interest to train machine learning models to decode different behavioural, physiological, and/or neural signals from film and vice versa. Given the extensive emotion annotations, a valuable avenue may be to determine how different behavioural features modulate the decoding accuracy of such models [55].

Finally, our multimodal dataset may also offer a useful resource for methods development, especially for fMRI data analysis. It is important to validate new methods on diverse datasets and problems can emerge if the neuroscience community relies overly on a small number of large datasets. A particular strength of the Emo-FilM dataset is the wealth of continuous behavioural annotations available that can be put in relationship with physiological and fMRI recordings.

In sum, we hope that the Emo-FilM dataset will contribute both to a deeper understanding of human emotion processing, and to enhanced training and validation of future tools to automatically extract and characterise biologically relevant information pertaining to emotion experience.

## Acknowledgments

We thank the staff of the BBL for technical support and advice, especially Bruno Bonet, Damien Marie, Frederic Grouiller. Oriane Marguet, Merlin Leuenberger and Mariane Brodier, students who helped with acquisition. This research was funded by the Swiss National Science Foundation (Sinergia grant CRSII5_180319) and benefited from resources of the Swiss Center of Affective Sciences at UNIGE.

## Author Contributions

EM: Conceptualization (supporting), Methodology (lead), Investigation (lead), Validation (supporting), Formal Analysis (lead), Software (equal), Supervision (supporting), Visualization (lead), Writing – Original Draft Preparation (lead), Writing – Review & Editing (lead)

SM: Supervision (supporting), Validation (lead), Writing – Original Draft Preparation (supporting), Writing – Review & Editing (lead)

LV: Methodology (supporting), Investigation (supporting), Writing – Review & Editing (supporting)

RF: Formal Analysis (supporting), Validation (supporting), Writing – Review & Editing (supporting)

MM: Software (equal), Writing – Review & Editing (supporting)

MP: Investigation (supporting), Writing – Review & Editing (supporting)

MAB: Investigation (supporting), Writing – Review & Editing (supporting)

PV: Funding Acquisition (equal), Conceptualization (lead), Supervision (supporting), Writing – Review & Editing (lead)

DvdV: Funding Acquisition(equal), Conceptualization (lead), Supervision (lead), Writing – Review & Editing (lead)

## Notes

doi:10.18112/openneuro.ds004892.v1.0.0

doi:10.18112/openneuro.ds004872.v1.0.0

### Competing Interest Statement

The authors have declared no competing interest.

